# The TFIIH subunits p44/p62 act as a damage sensor during nucleotide excision repair

**DOI:** 10.1101/643874

**Authors:** JT Barnett, J Kuper, W Koelmel, C Kisker, NM Kad

## Abstract

Nucleotide excision repair (NER) protects the genome following exposure to diverse types of DNA damage, including UV light and chemotherapeutics. Mutations in mammalian NER genes lead to diseases such as xeroderma pigmentosum, trichothiodystrophy, and Cockayne syndrome. In eukaryotes, the major transcription factor TFIIH is the central hub of NER. The core components of TFIIH include the helicases XPB, XPD, and five ‘structural’ subunits. Two of these structural TFIIH proteins, p44 and p62 remain relatively unstudied; p44 is known to regulate the helicase activity of XPD during NER whereas p62’s role is thought to be structural. However, a recent cryo-EM structure shows that p44, p62, and XPD make extensive contacts within TFIIH, with part of p62 occupying XPD’s DNA binding site. This observation implies a more extensive role in DNA repair beyond the structural integrity of TFIIH. Here, we show that p44 stimulates XPD’s ATPase but upon encountering DNA damage, further stimulation is only observed when p62 is part of the ternary complex; suggesting a role for the p44/p62 heterodimer in TFIIH’s mechanism of damage detection. Using single molecule imaging, we demonstrate that p44/p62 independently interacts with DNA; it is seen to diffuse, however, in the presence of UV-induced DNA lesions the complex stalls. Combined with the analysis of a recent cryo-EM structure we suggest that p44/p62 acts as a novel DNA-binding entity within TFIIH that is capable of recognizing DNA damage. This revises our understanding of TFIIH and prompts more extensive investigation into the core subunits for an active role during both DNA repair and transcription.

## Introduction

Cell survival relies on accurate replication and transcription of genetic material. When DNA is damaged, rapid, efficient repair is essential to ensure genome integrity. Defects in the nucleotide excision repair (NER) machinery lead to diseases such as x*eroderma pigmentosum* (XP), which is phenotypically characterized by UV hyper-sensitivity and increased incidence of skin cancers (1). Mutations found in many components of the NER pathway, including TFIIH can lead to XP. The core structure of TFIIH comprises two helicases; of these, XPB is enzymatically involved in both transcription and repair, whereas XPD’s activity is only essential for repair (2, 3). In addition, TFIIH possesses five other core subunits: p44, p62, p34, p8, and p52 (4-13). While it has been shown that p44 and p52 can stimulate XPD and XPB’s helicase activities respectively (8, 14, 15), most of the core subunits are relatively unstudied. A recent cryo-EM structure of TFIIH reveals significant interactions between p44, p62, and XPD (16). Part of p62 is located in XPD’s DNA binding site, suggesting p62 might assume a role in modulating XPD’s ability to interact with DNA. Although p62 is known to be important for TFIIH’s stability (17), and recruiting the complex to DNA (18, 19), p62’s major role is thought to be purely structural (20, 21).

Previous attempts to purify full-length p44 have been hampered by protein stability issues, however, a truncated form of p44 has been expressed and has been shown to activate the ATPase of XPD (3). In this study, we successfully purified full length p44 through co-expression with p62. We show that the complex formed by both proteins not only activates, but further accelerates XPD’s ATPase in the presence of damage. Remarkably, in the absence of XPD, we observe that p44/p62 independently binds to double-stranded DNA (dsDNA). Using single molecule fluorescence microscopy, we show that these newly defined DNA bound complexes randomly diffuse on non-damaged dsDNA, but are stalled in the presence of UV lesions, suggesting a direct response to damage. This hypothesis is supported by the in-depth analysis of a recent TFIIH cryo-EM structure (22), where p62 fragments were placed into unassigned regions of the EM maps. This suggests that p62 locates proximally to XPD’s DNA binding pocket when bound to single-stranded DNA (ssDNA). Our data thus reveal that p44/p62 represents a novel damage-specific DNA binding entity within TFIIH.

## Results & discussion

### p44/p62 co-purify as a complex that stimulate XPD’s ATPase activity

Both p44 and p62 make extensive contacts with XPD in TFIIH (16). To investigate this interaction, recombinant p44, p62, and XPD from *Chaetomium thermophilum* were utilized. All subunits share high sequence homology with the human TFIIH subunits and we have shown previously that they serve as ideal model for the human proteins (3, 23). Only the N-terminal region (residues 1-285) of p44, containing the von Willebrand domain (N-p44), has been successfully purified to date, nonetheless this is sufficient to activate XPD’s ATPase and helicase (3). By co-expressing full-length p44 with full-length p62 we successfully purified a tightly associated complex of p44/p62 that also interacts with XPD (**Figure 1**), thus providing the first successful purification of full length p44, suggesting that p44 requires p62 to fold correctly.

**Figure 1.**
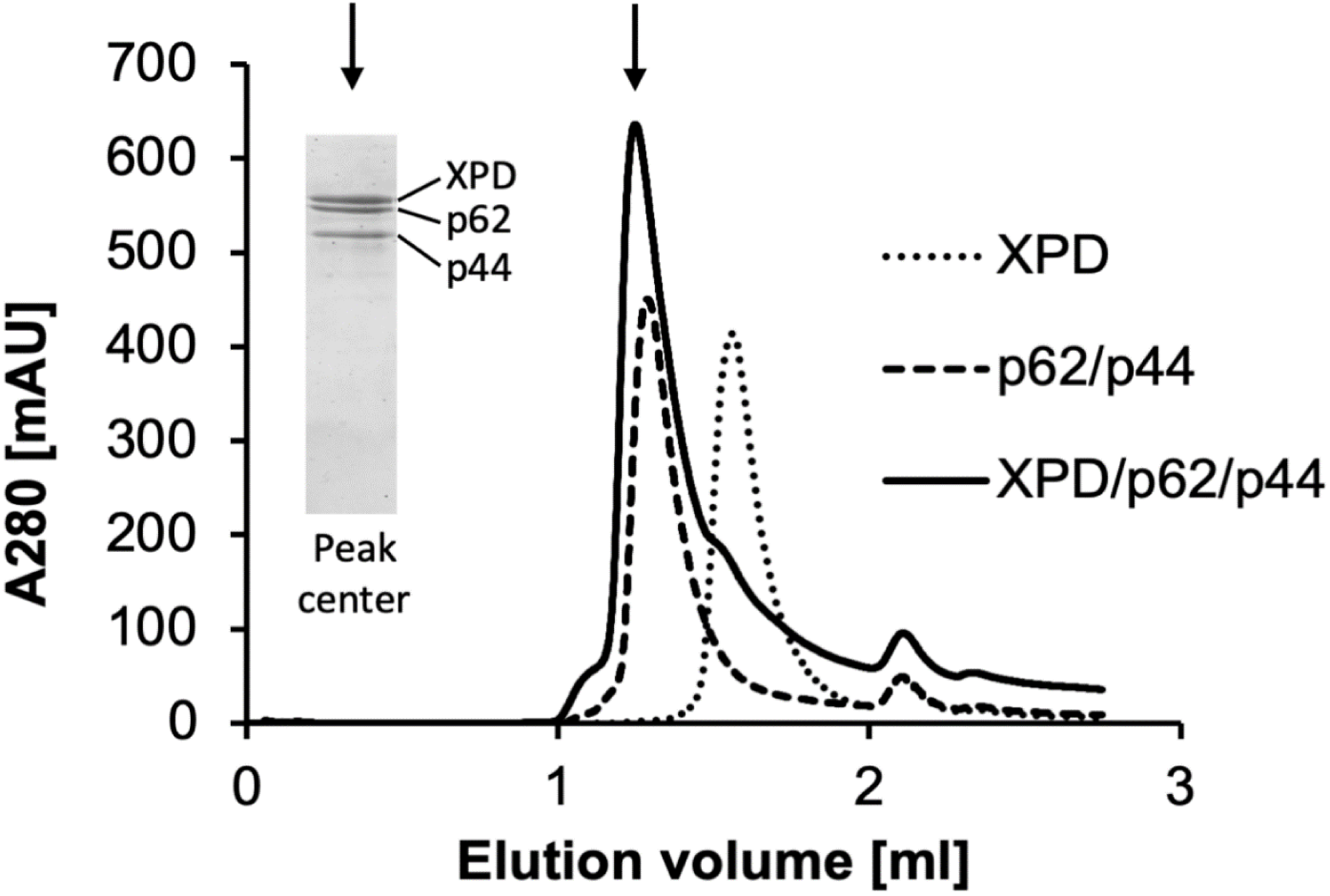
Size exclusion chromatography (SEC) analysis of XPD, the dimeric p44/p62 and the ternary XPD/p44/p62 complexes. SEC chromatograms of XPD (92 kDa), p44/p62 (132 kDa), and the XPD/p44/p62 (224 kDa) complex were performed in 20 mM HEPES pH 7.5, 250 mM NaCl, and 1 mM TCEP on a Superdex200 Increase 3.2/300. XPD was mixed with p44/p62 at an equal molar ratio (final concentration of 5 µM) prior to SEC. The inset shows the SDS-PAGE analysis of the center fraction from the XPD/p44/p62 elution.

To investigate the impact of the p44/p62 complex on XPD, we studied XPD’s ATPase activity in the presence of undamaged dsDNA or ssDNA substrates. In the presence or absence of DNA, XPD by itself possesses a slow ATPase (∼0.04 s^−1^). However, the addition of N-p44 significantly activates XPD’s ATPase on both dsDNA and ssDNA (0.136 s^−1^ and 0.504 s^−1^ respectively, p < 0.05 (**Figure 2A, B**)), consistent with previous reports (3). No further acceleration of XPD’s ATPase was observed in the presence of the full length p44/p62 complex with either undamaged dsDNA or ssDNA. However, remarkably, when a fluorescein moiety (recognized as damage by NER (24)) was introduced into a dsDNA substrate, we observed a further two-fold acceleration of XPD’s ATPase in the presence of p44/p62 compared to undamaged dsDNA (**Figure 2A, B)**. This was in contrast to N-p44 alone, which did not further accelerate XPD’s ATPase in the presence of damage, indicating that the ternary p44/p62:XPD complex is responsible for this damage-induced ATPase enhancement, and provides the first evidence that p44/p62 may assume a role in lesion detection. Such behavior is supported by the sensitization to UV irradiation observed in truncations of the yeast p62 homologue (17, 25, 26). To further investigate the XPD-activating role of p44 and p62, we analyzed XPD’s helicase activity on an open fork substrate (**Figure 2C**). Although no damage is present in the open fork substrate, p44/p62 significantly enhances XPD’s ability to successfully unwind the DNA substrate (two-fold more than XPD with N-p44), despite no observable change in ATPase activity, thus indicating that p44/p62 enhances the efficiency of the helicase activity.

**Figure 2.**
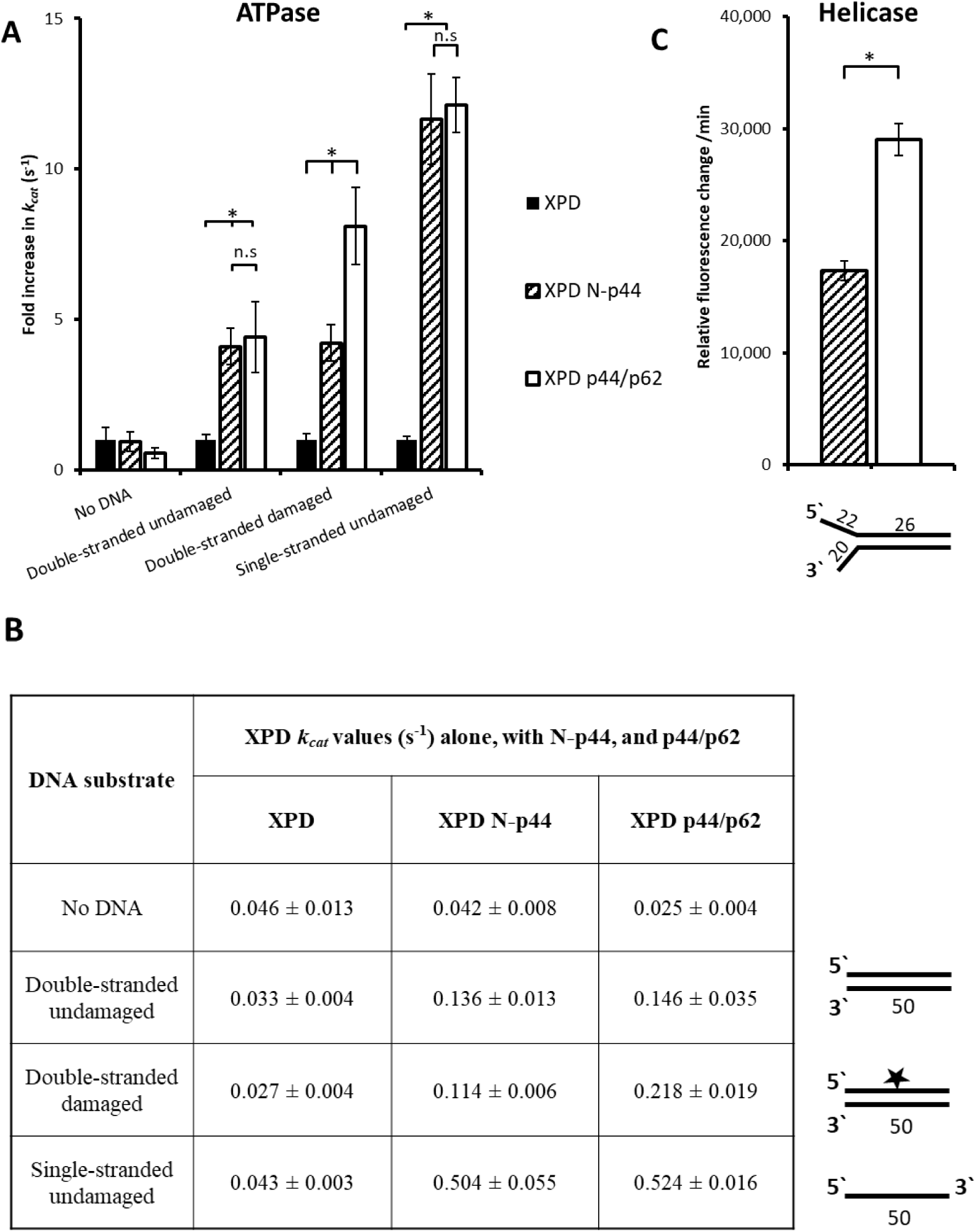
Steady-state ATPase and helicase activity of XPD in the presence of various DNA substrates and core TFIIH proteins. **(A)** The activity of XPD’s ATPase is stimulated by both N-p44 (dashed) and p44/p62 (white) on various DNA substrates. Values are k_cat_ given as a fold change compared to XPD alone (black). Errors are shown as S.E.M from 3 repeats. **(B)** Table of average k_cat_ values from (A) with a graphic of the DNA substrate. Length of substrate is given in nucleotides. **(C)** XPD’s helicase activity is stimulated by N-p44 (dashed) and p44/p62 (white) on an open fork substrate with lengths indicated in nucleotides. XPD alone displays no helicase activity (3). Errors are shown as S.E.M from 9 repeats. Statistical significance determined using a student’s t-test where * = p < 0.05, n.s = not statistically significant.

### The p44/p62 complex independently diffuses on DNA

To directly determine if p44/p62 interacts with DNA, we imaged individual fluorescently-tagged proteins binding to DNA tightropes in real time using fluorescence microscopy (27). In this assay, p44/p62 is fluorescently labelled using a quantum dot (Qdot) conjugated to the protein via its poly-histidine purification tag and an anti-His IgG antibody (28). Individual protein molecules are then imaged interacting with single DNA molecules in real time (**Figure 3A**). We observed substantial binding of p44/p62 complexes to dsDNA, and of these ∼80% diffused randomly (n = 599 total). To investigate the mechanism of p44/p62’s diffusion on the DNA, the diffusion constant and exponent (used to determine if motion is directionally biased) was calculated in high and low salt conditions (100 mM vs 10 mM KCl). At elevated KCl concentrations fewer p44/p62 complexes bound to DNA, and the calculated diffusion constant using mean-squared displacement analysis (27) showed no significant change (p > 0.05) between salt conditions (10 mM KCl 0.067 µm^2^/s ± 0.006 vs 100 mM KCl 0.042 µm^2^/s ± 0.010; **Figure 3B**), consistent with p44/p62 complexes sliding along the DNA helix (29). The diffusive exponent was found to be ∼0.9 in either salt condition, which indicates that there is no directional bias to the diffusive motion (30). Based on the estimated size of a p44/p62 complex conjugated to a Qdot, the diffusion constant is consistent with rotation-coupled diffusion around the backbone of the DNA helix (27, 31, 32). Altogether, these data provide the first observation that the p44/p62 complex can bind DNA and randomly diffuse, independently of XPD.

**Figure 3.**
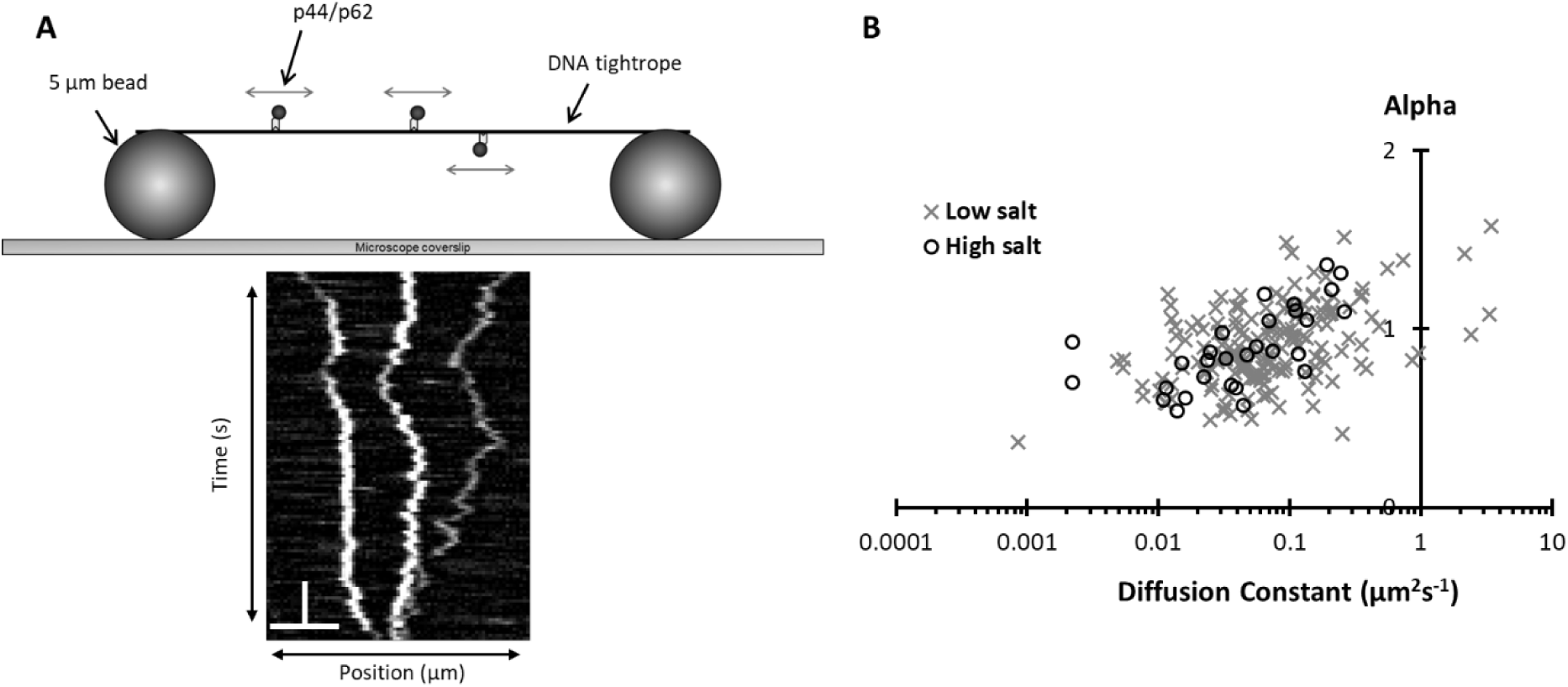
Imaging of p44/p62’s motility on DNA tightropes. **(A)** Schematic drawing of the DNA tightrope assay: Qdot-labelled proteins are imaged interacting with single molecules of DNA suspended between 5 µm diameter beads. The motion of three diffusing p44/p62 complexes are transformed into a kymograph by plotting their position along the DNA against time. The scale-bar represents 10 s and 10 µm. **(B)** Diffusion constant versus exponent (alpha) at high (circles) and low salt (crosses). The average diffusion constant values are given in the main text. Average alpha exponent values were 0.89 ±0.04 and 0.91 ±0.02 in high and low salt respectively.

### p44/p62 complexes are stalled by the presence of damaged DNA

Recently Greber et al., (2019) showed by cryo-EM, that a portion of p62 is located in XPD’s DNA binding site within the DNA-free transcriptional apo-TFIIH structure (16). However, when XPD interacts with DNA within the NER pathway it is conceivable that p62 undergoes a conformational change and is thereby able to interact directly with the DNA. This could offer a putative mechanism by which XPD’s ATPase is stimulated by p44/p62 in the presence of damage. To investigate this hypothesis, we determined p44/p62’s affinity for different DNA substrates relevant to NER by fluorescence polarization (**Figure 4A**). Complementary to our observations with DNA tightropes, p44/p62 was found to bind dsDNA with a *K*_*D*_ of 1.18 µM. Interestingly, DNA intermediates expected to form during NER showed stronger binding by p44/p62 than dsDNA (*K*_*D*_ (ssDNA) 0.84 µM and *K*_*D*_ (open fork) 0.21 µM). The tighter binding to these NER intermediates strengthens the suggestion that p44/p62 may contact DNA within TFIIH.

**Figure 4.**
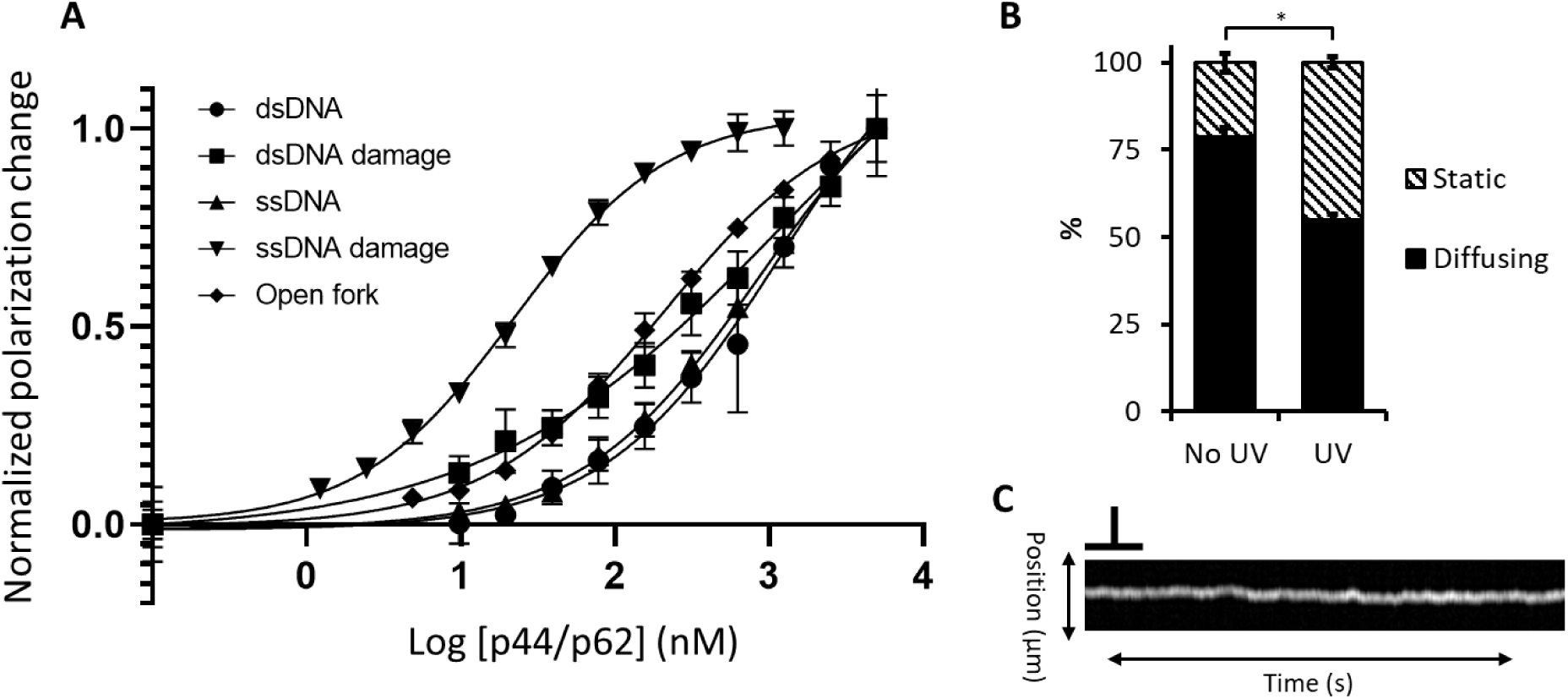
p44/p62 binding affinities for different DNA substrates. **(A)** Fluorescence polarization was used to determine the binding of various substrate to increasing concentrations of p44/p62. p44/p62 displayed an intermediate affinity for dsDNA irrespective of the presence of a fluorescein DNA damage, but a higher affinity for open fork and ssDNA. The tightest binding was observed with ssDNA containing fluorescein DNA damage. Data were plotted and fitted in GraphPad, presented here as normalized values with error bars representing the SD from at least three repeats. **(B)** Single molecule fluorescence imaging reveals that the number of diffusing p44/p62 complexes on dsDNA tightropes decreases in the presence of UV-induced DNA damage. Data are average percentages from 8 (No UV) or 6 (UV) experimental repeats with the S.E.M as errors bars, Statistical significance determined using a student’s t-test where * = p < 0.05. **(C)** Example kymograph of a molecule showing constrained diffusion. The scale-bar represents 10 s and 1 µm.

Since p44/p62 further accelerated XPD’s ATPase specifically on damaged DNA we also investigated how the addition of damage affected the affinity of p44/p62 for different DNA damage containing substrates. p44/p62 bound dsDNA containing a fluorescein moiety with similar affinity to undamaged DNA (*K*_*D*_ 1.23 µM and 1.18 µM respectively). In contrast, we observed a striking 40-fold increase in binding affinity to a ssDNA substrate containing a central fluorescein lesion (*K*_*D*_ 0.02 µM) compared to the ssDNA substrate without damage.

To further analyze these findings, we investigated the effect of DNA damage on the behavior of single p44/p62 complexes on DNA tightropes irradiated with 500 J/m^2^ of 254 nm UV light, which generates cyclobutane pyrimidine dimers and 6-4 photoproducts (33). We observed a striking and significant decrease in the number of diffusing p44/p62 complexes from ∼80% to 50% ((p < 0.05) **Figure 4B**). These results clearly indicate that p44/p62 independently recognizes DNA damage and stalls at damaged sites. We also observed a change in the mechanism of p44/p62’s diffusion in the presence of UV damage; a population of p44/p62 complexes now appeared limited in their total excursion distance on DNA (**Figure 4C**), similar to the damage recognition protein XPC, which undergoes constrained diffusion around a lesion (34, 35). Although p44/p62 cannot discriminate DNA damage in a dsDNA substrate without XPD (**Figure 4A**), the observed stalling in the DNA tightrope assay suggests that p44/p62 may have located exposed ssDNA regions containing UV lesions. Together these data implicate a role for p44/p62 towards recognizing DNA lesions within TFIIH.

### The structural basis of p62-DNA contacts within TFIIH

The recent cryo-EM structure of TFIIH bound to DNA shows the path of single-stranded DNA as it is passed from XPB to XPD (22). In this structure, p44 is located in the same position as in the apo-TFIIH(16), but p62 is unassigned. However, additional electron densities can be observed in the structure at a position similar to p62 in apo-TFIIH (**Figure 5A**) (16). We modelled regions of p62 from the apo-TFIIH into the DNA-bound TFIIH structure and found excellent agreement with the position of p62 (**Figure 5B**). This model suggests that the electron density close to XPD could represent N-terminal regions of p62. Closer examination of the DNA binding groove within XPD shows that p62 occupies this site in the absence of DNA (**Figure 5C**), i.e. in the apo TFIIH structure. Analysis of the same region in the DNA-bound structure suggests that the p62 loop could undergo a conformational change to form a helical bundle that interacts with the Arch domain of XPD and the ssDNA bound to XPD (**Figure 5D**).

**Figure 5.**
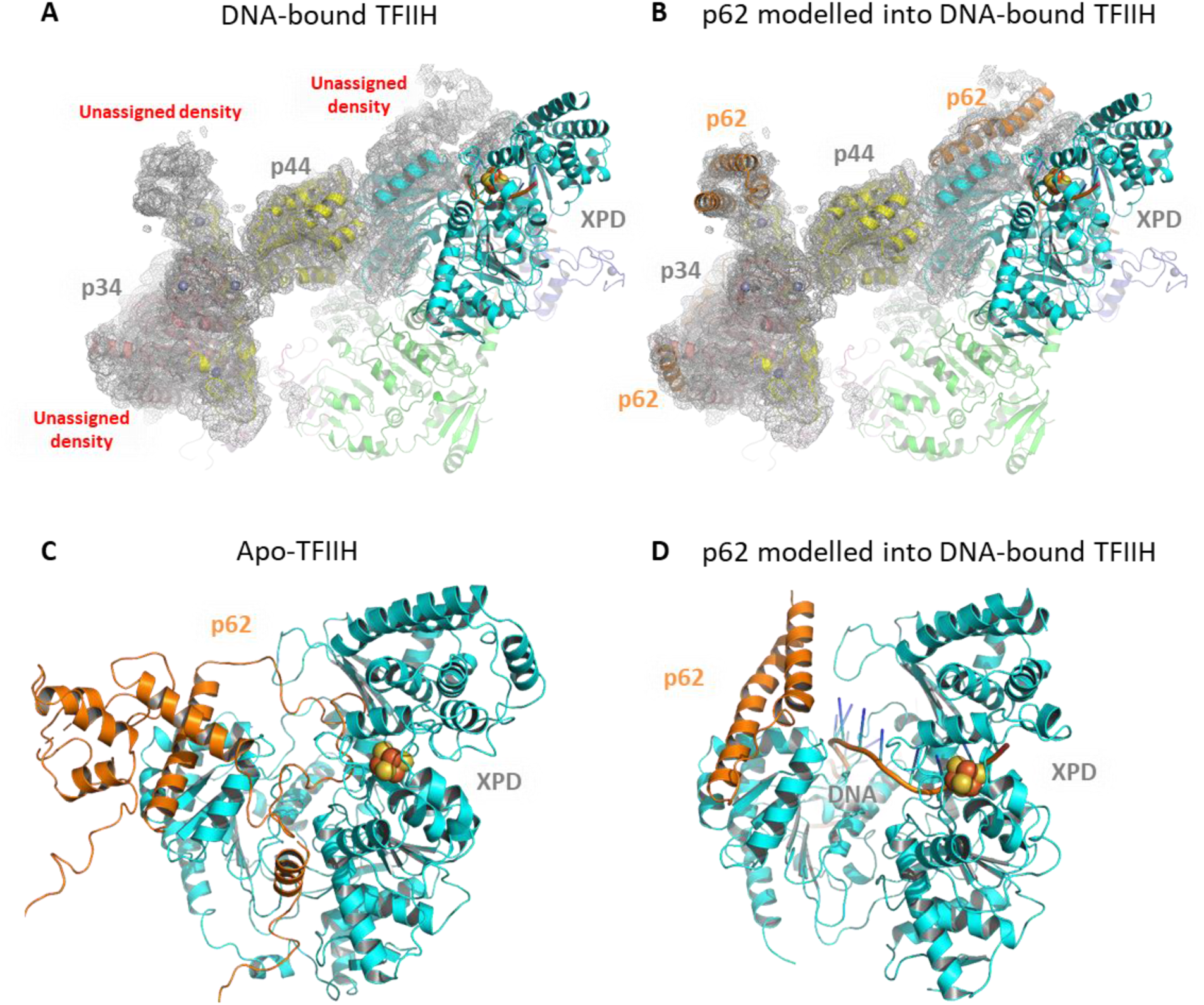
The position of p62 in apo- and DNA bound TFIIH structures. **(A)** Representation of the Kokic et al DNA-bound TFIIH structure (PDB:6RO4) shown in cartoon mode with the EM-envelope displayed as grey mesh. **(B)**The unmodelled density around p44/p34 was fit using p62 from the Greber et al apo-TFIIH structure and the 3 helical bundle (residues 450-547) superimposed with an rmsd of 0.5 Å after rebuilding. Residues 397-450 differ in orientation due the concerted movement of XPD-p44 in 6ro4. The additional part of p62 interacting with XPD was modelled according to the density with no defined amino acid sequence. **(C)** Close-up view of XPD and p62 based on the apo-TFIIH structure (PDB:6NMI), with the iron sulphur cluster in XPD shown as spheres. **(D)** Equivalent view of XPD from the DNA-bound TFIIH structure with the additional electron density modelled as p62.

In conclusion, we present the first mechanistic characterisation of the non-helicase TFIIH subunits p44/p62. Our biochemical data clearly show that the p44/p62 complex binds to DNA but displays a profoundly tighter affinity for ssDNA containing damage (**Figure 4**). Modelling of non-assigned densities in a recent cryo-EM structure indicate p62 to be in a position where it could interact with XPD and DNA in a substrate dependent manner (**Figure 5**). Taken together, these data suggest that p44/p62 interacts with ssDNA bound to XPD and is capable of modulating its helicase/ATPase activity through interactions with the helicase and/or Arch domains. This provides a structural explanation for the observed DNA damage-dependent increase in XPD’s ATP turnover in the presence of p44/p62 (**Figure 2**). Furthermore, we have used single molecule imaging to directly demonstrate that p44/p62 complexes are capable of recognizing UV damage and consequently stalling on DNA (**Figures 3 & 4**). This behavior could be important for stalling XPD at a lesion and triggering damage verification. Nonetheless, our results clearly show that the p44/p62 complex plays an active role in TFIIH’s recognition of DNA damage and indicates p44/p62 can no longer be thought of simply as ‘structural subunits’.

## Methods

### Purification

The genes encoding p44 and p62 were cloned from *C. thermophilum* cDNA. p62 was cloned into the pETM-11 vector (EMBL) without a tag. p44 was cloned into the pBADM-11 vector (EMBL) containing an N-terminal hexa-Histidine tag followed by a TEV cleavage site. p62 and p44 were co-expressed in *E. coli* BL21 CodonPlus (DE3) RIL cells (Agilent) and were co-purified via immobilized metal affinity chromatography (Ni TED, Machery-Nagel), followed by size exclusion chromatography (SEC), and anion exchange chromatography (AEC). SEC was conducted with a HiLoad 16/600 Superdex 200 prep grade column (GE Healthcare) in 20 mM Hepes pH 7.5, 250 mM NaCl, and 1 mM TCEP. AEC was conducted with a MonoQ 5/50 GL column (GE Healthcare). The proteins were eluted via a salt gradient ranging from 50 to 1000 mM NaCl. AEC buffers were composed of 20 mM HEPES pH 7.5, 50/1000 mM NaCl, and 1 mM TCEP. The p62/p44 protein complex was concentrated to approximately 20 mg/ml and flash frozen in liquid nitrogen for storage.

XPD and N-p44 (1-285) from *C. thermophilum* were cloned as described previously (3). XPD was expressed as N-terminally His-tagged proteins in *E. coli* ArcticExpress (DE3)-RIL cells (Agilent). Cells were grown in TB medium at 37°C until they reached an OD_600_ of 0.6-0.8. Expression was initiated by the addition of 0.05% L-arabinose and performed at 11°C for 20 h. N-p44 was expressed as N-terminally His-tagged protein in *E. coli* BL21-CodonPlus (DE3)-RIL cells (Stratagene). Cells were grown as described for ctXPD and expression was started by adding 0.1 mM IPTG at 14°C for 18 h. XPD and N-p44 were purified to homogeneity by metal affinity chromatography (Ni-IDA, Macherey&Nagel) as described previously (3) followed by size exclusion chromatography (SEC) (20 mM HEPES pH 7.5, 200 mM NaCl) and an additional anion exchange chromatography (AEC) step in the case of XPD. AEC was performed using a MonoQ 5/50 GL column (GE Healthcare) with 20 mM HEPES pH 7.5, 50 mM NaCl, and 1 mM TCEP as loading buffer and the same buffer containing 1 M NaCl was used for elution. The final buffer after AEC was 20 mM HEPES pH 7.5, 250 mM NaCl, and 1 mM TCEP. The proteins were concentrated to at least 5 mg/ml based on their calculated extinction coefficient using ProtParam (SwissProt) and then flash frozen for storage at -80°C.

### Size exclusion chromatography

Size exclusion chromatography of XPD, p44/p62, and the XPD/p44/p62 complex was performed in 20 mM HEPES pH 7.5, 250 mM NaCl, 1 mM TCEP at a concentration of 5 µM. XPD was mixed with p44/p62 at an equal molar ratio with the final concentration of 5 µM prior to the size exclusion chromatography analysis. Size exclusion chromatography was performed on an Äkta Pure system using a Superdex 200 5/150GL at a flow rate of 0.3 ml/min.

### Fluorescence anisotropy

DNA binding was analyzed by fluorescence anisotropy employing the DNA substrates indicated in the methods section for the ATPase with the addition of a Cy3 label on the 3’ end F26,50 with and without the modification. In addition, we used a hairpin DNA representing the open fork structure with a Cy3 label on the 3’ end (5’TTT TTT TTT TTT TTTCCC GGC CAT GC *GAA* GC ATG GCC GGG TTT TT3’). Assays were carried out in 20 mM HEPES pH 7.5, 30 mM KCl, 5 mM MgCl_2_, 1 mM TCEP, and 5 nM DNA at room temperature. The protein was used at concentrations of 2 to 5000 nM as indicated. Fluorescence was detected at an excitation wavelength of 540 nm and an emission wavelength of 590 nm with a Clariostar plate reader (BMG labtech). The gain was adjusted to a well containing buffer and DNA but no protein. Curves were fitted with GraphPad Prism and represent the averages of at least three different reactions.

### ATPase assay

dsDNA substrates used:

F26,50 contains a fluorescein moiety covalently attached to thymine (*);

5’GACTACGTACTGTTACGGCTCCATCT*CTACCGCAATCAGGCCAGATCTGC 3’

The reverse complementary sequence to F26,50;

5’GCAGATCTGGCCTGATTGCGGTAGCGATGGAGCCGTAACAGTACGTAGTC 3’

F26,50 without the fluorescein moiety;

5’GACTACGTACTGTTACGGCTCCATCTCTACCGCAATCAGGCCAGATCTGC 3’

The NADH-coupled ATPase assay was performed as described previously (33) in plate reader format. Imaging buffer containing the NADH-reaction components was supplemented with 1 mM fresh TCEP, protein (100 nM (equimolar concentrations for XPD N-p44 and XPD p44/p62)), and 50 nM of DNA substrate. The reaction was started with the addition of 1 mM ATP to each well, and the change in OD340 (NADH) was monitored every 8 seconds/well over 30 minutes at room temperature in a Clariostar plate reader. The rates of NADH consumption were used to calculate *k*_*cat*_. Reactions were repeated 3 times, and S.E.M used as errors values.

### *In vitro* helicase assay

Helicase activity was analyzed utilizing a fluorescence-based assay. We used an open fork substrate with a Cy3 label at the 3’ end of the translocated strand where unwinding of the DNA substrate reduces quenching of the Cy3 fluorescence.

5’AGCTACCATGCCTGCACGAATTAAGCAATTCGTAATCATGGTCATAGC-Cy3 3’

and a dabcyl modification on the 5’ end of the opposite strand

5’ Dabcyl-GCTATGACCATGATTACGAATTGCTTGGAATCCTGACGAACTGTAG 3’

Assays were carried out in 20 mM HEPES pH 7.5, 50 mM KCl, 5 mM MgCl_2_, and 1 mM TCEP. DNA was used at a concentration of 250 nM. Helicase activity was measured with equimolar concentrations of XPD, p44, and/or p62. The mix of reagents, were preincubated at 37°C and the reaction was subsequently started with the addition of 5 mM ATP. Kinetics were recorded with a Fluostar Optima plate reader (BMG labtech). Fluorescence was detected at an excitation wavelength of 550 nm (slit width, 2 nm) and an emission wavelength of 570 nm (slit width, 2 nm). Initial velocities were fitted with the MARS software package (BMG labtech) and represent the averages of at least three different reactions and two independent protein batches.

### Single Molecule DNA Tightrope Assay

For a detailed protocol see (28). p44/p62 interactions with DNA were studied in imaging buffer (20 mM Tris pH 8.0, 10 mM KCl (100 mM for high salt), 5 mM MgCl_2_, 1 mM TCEP). Videos for diffusion analysis were collected between 30 seconds and 5 minutes at 10 frames per second. Video analysis was performed in ImageJ as described previously (27). Experiments using UV damaged lambda are described elsewhere (33). Briefly, lambda DNA was irradiated to 500 J/m^2^ using a 254 nm lamp immediately before constructing DNA tightropes. Exposure of DNA to this wavelength randomly induces CPD and (6-4) Photoproduct lesions that are recognised as damage by NER (24, 36).

### Structural Model Building

The additional p62 fragments were built into the cryo-EM map of entry 6ro4 using manual model building features in coot and coot real space refinement. For the C-terminal part, the corresponding segments of p62 from the PDB entry 6nmi (residues 397-547) were used and fitted into the density. The N-terminal helical bundle was built *de novo*.

## Data availability

All data will be made available through the University of Kent academic repository (https://kar.kent.ac.uk/).

## Acknowledgements

We would like to thank the members of the Kad group for useful discussions. This work was supported by the Biotechnology and Biological Sciences Research Council BB/P00847X/1, BB/M019144/1, BB/I003460/1 to NMK and BB/M01603X/1 to NMK and JTB and by the German Research Foundation KI-562/7-1 to CK. The authors declare no conflict of interest.

## Author contributions

Collected data: JTB, JK, WK. Designed experiments: JTB, JK, CK, NMK. Analysed Data: JTB, JK, NMK. Wrote paper: JTB, JK, CK, NMK.

## References

1. Bradford PT, et al. (2011) Cancer and neurologic degeneration in xeroderma pigmentosum: long term follow-up characterises the role of DNA repair. J Med Genet 48(3):168–176.

2. Coin F, Oksenych V, & Egly JM (2007) Distinct roles for the XPB/p52 and XPD/p44 subcomplexes of TFIIH in damaged DNA opening during nucleotide excision repair. Mol Cell 26(2):245–256.

3. Kuper J, et al. (2014) In TFIIH, XPD helicase is exclusively devoted to DNA repair. PLoS biology 12(9):e1001954.

4. Roy R, et al. (1994) The DNA-dependent ATPase activity associated with the class II basic transcription factor BTF2/TFIIH. J Biol Chem 269(13):9826–9832.

5. Greber BJ, et al. (2017) The cryo-electron microscopy structure of human transcription factor IIH. Nature 549(7672):414.

6. Sung P, et al. (1993) Human xeroderma pigmentosum group D gene encodes a DMA helicase. Nature 365:852.

7. Schultz P, et al. (2000) Molecular structure of human TFIIH. Cell 102(5):599–607.

8. Luo J, et al. (2015) Architecture of the Human and Yeast General Transcription and DNA Repair Factor TFIIH. Mol Cell 59(5):794–806.

9. Araujo SJ, et al. (2000) Nucleotide excision repair of DNA with recombinant human proteins: definition of the minimal set of factors, active forms of TFIIH, and modulation by CAK. Genes Dev 14(3):349–359.

10. Svejstrup JQ, et al. (1995) Different forms of TFIIH for transcription and DNA repair: Holo-TFIIH and a nucleotide excision repairosome. Cell 80(1):21–28.

11. Habraken Y, Sung P, Prakash S, & Prakash L (1996) Transcription factor TFIIH and DNA endonuclease Rad2 constitute yeast nucleotide excision repair factor 3: implications for nucleotide excision repair and Cockayne syndrome. Proceedings of the National Academy of Sciences 93(20):10718–10722.

12. Sung P, Prakash L, Matson SW, & Prakash S (1987) RAD3 protein of Saccharomyces cerevisiae is a DNA helicase. Proceedings of the National Academy of Sciences 84(24):8951–8955.

13. Sung P, Prakash L, Weber S, & Prakash S (1987) The RAD3 gene of Saccharomyces cerevisiae encodes a DNA-dependent ATPase. Proceedings of the National Academy of Sciences 84(17):6045–6049.

14. Jawhari A, et al. (2002) p52 Mediates XPB function within the transcription/repair factor TFIIH. J Biol Chem 277(35):31761–31767.

15. Coin F, et al. (1998) Mutations in the XPD helicase gene result in XP and TTD phenotypes, preventing interaction between XPD and the p44 subunit of TFIIH. Nature Genetics 20(2):184–188.

16. Greber BJ, Toso DB, Fang J, & Nogales E (2019) The complete structure of the human TFIIH core complex. Elife 8.

17. Gervais V, et al. (2004) TFIIH contains a PH domain involved in DNA nucleotide excision repair. Nat Struct Mol Biol 11(7):616–622.

18. Yokoi M, et al. (2000) The xeroderma pigmentosum group C protein complex XPC-HR23B plays an important role in the recruitment of transcription factor IIH to damaged DNA. Journal of Biological Chemistry 275(13):9870–9875.

19. Sugasawa K, et al. (1996) HHR23B, a human Rad23 homolog, stimulates XPC protein in nucleotide excision repair in vitro. Molecular and cellular biology 16(9):4852–4861.

20. Jawhari A, et al. (2004) Domain architecture of the p62 subunit from the human transcription/repair factor TFIIH deduced by limited proteolysis and mass spectrometry analysis. Biochemistry 43(45):14420–14430.

21. Di Lello P, et al. (2006) Structure of the Tfb1/p53 complex: Insights into the interaction between the p62/Tfb1 subunit of TFIIH and the activation domain of p53. Molecular cell 22(6):731–740.

22. Kokic G, et al. (2019) Structural basis of TFIIH activation for nucleotide excision repair. Nat Commun 10(1):2885.

23. Radu L, et al. (2017) The intricate network between the p34 and p44 subunits is central to the activity of the transcription/DNA repair factor TFIIH. Nucleic Acids Res.

24. Buechner CN, et al. (2014) Strand-specific recognition of DNA damages by XPD provides insights into nucleotide excision repair substrate versatility. J Biol Chem 289(6):3613–3624.

25. Gileadi O, Feaver WJ, & Kornberg RD (1992) Cloning of a subunit of yeast RNA polymerase II transcription factor b and CTD kinase. Science 257(5075):1389.

26. Matsui P, DePaulo J, & Buratowski S (1995) An interaction between the Tfb1 and Ssl1 subunits of yeast TFIIH correlates with DNA repair activity. Nucleic Acids Research 23(5):767–772.

27. Kad NM, Wang H, Kennedy GG, Warshaw DM, & Van Houten B (2010) Collaborative dynamic DNA scanning by nucleotide excision repair proteins investigated by single-molecule imaging of quantum-dot-labeled proteins. Mol Cell 37(5):702–713.

28. Springall L, Inchingolo AV, & Kad NM (2016) DNA-Protein Interactions Studied Directly Using Single Molecule Fluorescence Imaging of Quantum Dot Tagged Proteins Moving on DNA Tightropes. Methods Mol Biol 1431:141–150.

29. von Hippel PH & Berg OG (1989) Facilitated target location in biological systems. J Biol Chem 264(2):675–678.

30. Saxton MJ (2001) Anomalous subdiffusion in fluorescence photobleaching recovery: a Monte Carlo study. Biophys J 81(4):2226–2240.

31. Schurr JM (1979) The one-dimensional diffusion coefficient of proteins absorbed on DNA. Biophysical Chemistry 9(4):413–414.

32. Hughes CD, et al. (2013) Real-time single-molecule imaging reveals a direct interaction between UvrC and UvrB on DNA tightropes. Nucleic Acids Res 41(9):4901–4912.

33. Barnett JT & Kad NM (2019) Understanding the coupling between DNA damage detection and UvrA’s ATPase using bulk and single molecule kinetics. FASEB journal : official publication of the Federation of American Societies for Experimental Biology 33(1):763–769.

34. Kong M, et al. (2016) Single-molecule imaging reveals that Rad4 employs a dynamic DNA damage recognition process. Molecular cell 64(2):376–387.

35. Cheon NY, Kim H-S, Yeo J-E, Schärer OD, & Lee JY (2019) Single-molecule visualization reveals the damage search mechanism for the human NER protein XPC-RAD23B. Nucleic acids research.

36. Sancar A & Rupp WD (1983) A novel repair enzyme: UVRABC excision nuclease of Escherichia coli cuts a DNA strand on both sides of the damaged region. Cell 33(1):249–260.

